# Transitioning to confined spaces impacts bacterial swimming and escape response

**DOI:** 10.1101/2021.09.15.460467

**Authors:** Jonathan B. Lynch, Nicholas James, Margaret McFall-Ngai, Edward G. Ruby, Sangwoo Shin, Daisuke Takagi

## Abstract

Symbiotic bacteria often navigate complex environments before colonizing privileged sites in their host organism. Chemical gradients are known to facilitate directional taxis of these bacteria, guiding them towards their eventual destination. However, less is known about the role of physical features in shaping the path the bacteria take and defining how they traverse a given space. The flagellated marine bacterium *Vibrio fischeri*,which forms a binary symbiosis with the Hawaiian bobtail squid, *Euprymna scolopes*, must navigate tight physical confinement, squeezing through a bottleneck constricting to ~2 μm in width on the way to its eventual home. Using microfluidic *in vitro* experiments, we discovered that *V. fischeri* cells alter their behavior upon entry into confined space, straightening their swimming paths and promoting escape from confinement. Using a computational model, we attributed this escape response to two factors: reduced directional fluctuation and a refractory period between reversals. Additional experiments in asymmetric capillary tubes confirmed that *V. fischeri* quickly escape from tapered ends, even when drawn into the ends by chemoattraction. This avoidance was apparent down to a limit of confinement approaching the diameter of the cell itself, resulting in a balance between chemoattraction and evasion of physical confinement. Our findings demonstrate that non-trivial distributions of swimming bacteria can emerge from simple physical gradients in the level of confinement. Tight spaces may serve as an additional, crucial cue for bacteria while they navigate complex environments to enter specific habitats.

**Significance Statement:** Symbiotic bacteria that navigate to and through specific host tissues often face tight physical confinement. This work reveals that confinement-associated changes in swimming can dramatically alter taxis, shaping bacterial localization in conjuncture with other motility-directing cues. This work helps explain how bacteria can avoid getting stuck in confined areas while transiting to privileged spaces, adding confinement as an environmental cue that symbiotic bacteria use to shape their motility behavior.

## Introduction

Motility is essential to the physiology of many bacteria. One of the most common modes of bacterial motility is coordinated rotation of extended structures called flagella, which utilize physical anisotropy to propel cells through viscous fluids (1, 2). Coordination of flagellar rotation results in directional movement and, when combined with detection of physical and chemical cues, enables bacteria to migrate to or from particular sites, facilitating activities such as predation (3), accessing preferred niches (4, 5), and pathogenesis (6). The directionality of flagellar swimming has been classically described in *Escherichia coli* by the “run-and-tumble” model (7), but flagella are also involved in other motility strategies, such as flicking (8, 9) and wrapping (10, 11). These swimming modalities each involve different specific flagellar actions, and their effects on cellular behavior depend on physical characteristics of the cell’s environment, with seemingly straightforward changes in environment drastically shaping overall bacterial behavior (12).

Swimming and reorientation in bulk fluid by flagellated bacteria are behaviors that have been well-studied (13), but recent findings have highlighted that these behaviors do not translate well to swimming in more complex environments. This may be due to changes in environmentally-stimulated flagellar organization or activity, or because of interactions between the cell and the physical environment that do not exist in bulk fluid. For example, physical forces cause flagellated swimmers to associate with surfaces (14, 15), which has interesting consequences in conditions of heterogeneous surface features (16). Relatedly, changes in the physical environment can also impact behaviors like chemotaxis, further highlighting the significance of physical features in processes like nutrient acquisition and habitat access (17).

Physical confinement is one important factor that bacteria commonly face during motility, and confinement effects can profoundly alter their trajectory and migration patterns (18); nevertheless, this factor is explicitly excluded from open-space swimming measurements. Even under moderate levels of confinement, swimming microorganisms have been observed to accumulate near a solid surface, produce circular trajectories, and scatter from obstacles (15, 19). Recent work revealed that even between bacteria with similar flagellar organizations, there are large variations in swimming tendencies, especially with regards to surface association and confined movement (19). The drastic changes in the behavior of confined bacteria as well as the underlying physical mechanisms are not well understood, and highlight that open space swimming characterizations will often not accurately generalize to swimming in other physical conditions. Understanding the impact that confinement has on flagellar swimming is especially important for host-associated bacteria, which often navigate tight spaces to reach privileged host sites (20).

The symbiotic marine bacterium *Vibrio (Aliivibrio) fischeri* offers a valuable model for studying the influence that transitioning to confined space has on flagellar swimming. Several strains of this bacterium transit from open ocean water to stably colonize animal hosts, such as the Hawaiian bobtail squid (*Euprymna scolopes*) (21). In the well characterized squid-vibrio symbiosis, *V. fischeri* traverses distinct microenvironments before reaching its permanent residence in the deep crypts of the juvenile squid light organ. The penultimate microenvironment *V. fischeri* reaches before the crypts --- the so-called bottleneck --- can close to widths of less than 2 microns (22, 23), preventing most cells from completing this journey. This situation means that, as *V. fischeri* colonizes the squid, it must transition from effectively no confinement to an extremely tight degree of confinement. As flagellar motility has previous been shown to be essential for its symbiotic organ colonization (5, 24), and as other *Vibrio spp*. use flagella to colonize similarly host tissues with some degree of confinement (25), we reasoned that *V. fischeri* would provide an excellent tool for examining changes in swimming behaviors during transitions between environments of varying confinement.

Here, we use microfluidics, high-speed microscopy, and mathematical modeling to describe the behavior of bacteria swimming into a confined space from an unconfined one. We find that while speed remains nearly constant across conditions, cells swimming under confined conditions tend to swim straighter than less confined cells. This adjustment in swimming directionality is paired with a delayed frequency of sharp reversals. Together, these new behaviors promote an exiting of highly confined spaces, an outcome that we observed through both simulation and experimental observation. We also found that this apparent aversion to confinement can be counterbalanced with chemoattraction, leading to a stable spatial organization of cells. These factors may explain a critical behavior used by bacteria while they navigate between free-living and symbiotic habitats.

## Results

### Physical confinement alters swimming behavior

*V. fischeri* displays a generally non-directional pattern of swimming in open space (Movie S1); however, due to previously described work finding that asymmetric physical environments promote spatially organized bacterial populations (26), we hypothesized that confinement similar to that seen during colonization would stimulate a more directed or organized swimming activity. To test the effects that physical confinement had on bacterial swimming behavior, we designed dead-end microfluidics devices with varying levels of two-dimensional confinement (Fig. 1A). These devices contained a central “open” chamber, in which there is effectively no confinement in any direction, as well as several perpendicular “confined” channels, in which there is little confinement in the x-y direction, but the bacteria are confined on the top and the bottom (z-direction) (Fig. 1A). Importantly, there was no directional background flow throughout the microfluidic apparatus (27).

**Figure 1.**
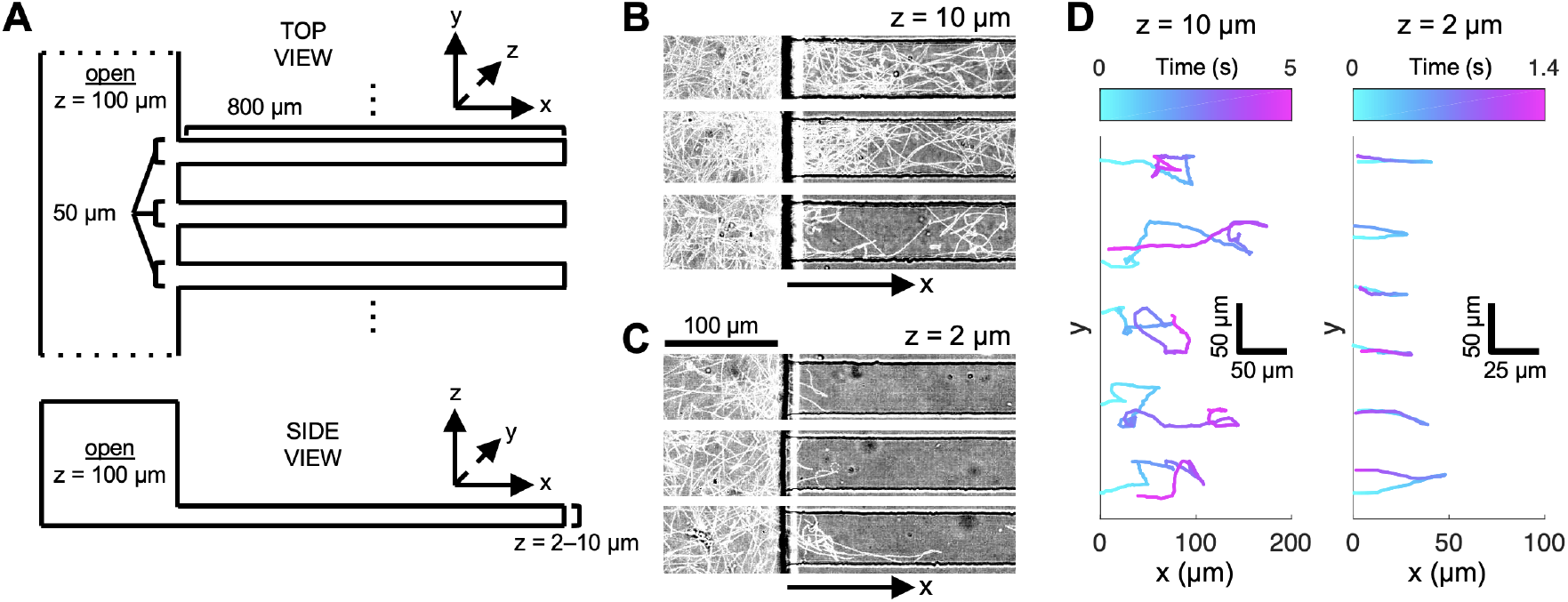
Confinement promotes channel escape in *V. fischeri*. A) Schematic of confinement channels for 2 and 10 μm confinement. B) Collapsed traces of *V. fischeri* swimming under 10μm z-confinement over 10 seconds. C) Collapsed traces of *V. fischeri* swimming under 2 μm z-confinement over 10 seconds. Left of vertical black lines is open chamber, right of black line is confined channel. D) Selected traces of *V. fischeri* cells entering noted z confinement. Color denotes time in seconds.

We measured swimming under varying confinement after loading these devices with a suspension of the well-studied *V. fischeri* isolate ES114 (28) in particle-free filter-sterilized ocean water. This medium choice was made to minimize consumable nutrients and the potential production of chemotactic gradients. Previous electron microscopy images showed that the comma-shaped cell bodies of intact *V. fischeri* cells are ~0.6-1 μm wide (29, 30), so to create relatively confined and unconfined environments, we used devices with two z heights, defining the two-sided confinement spacing: 10 μm, in which *V. fischeri* would be relatively less confined, and 2 μm, in which *V. fischeri* would be tightly confined. We were surprised to discover that *V. fischeri* cells displayed distinctive swimming behaviors as they moved from the open chamber into the differently confined channels (Movies S2, S3). Specifically, while cells entering the 10 μm channels swam freely throughout the channel (Fig. 1B), cells entering the tightly confined 2 μm channels quickly and efficiently exited the channels (Fig. 1C). These contrasting behaviors were even more marked when we tracked individual bacteria in each confinement (Fig. 1D).

### Confinement reduces residency

We tracked >300 individual trajectories of *V. fischeri* cells entering either confined or unconfined channels, and quantified differences in swimming parameters between cells entering these two differently confined channels. *V. fischeri* moving into confined channels penetrate more shallowly (Fig. 2A) and spend less time in the channels (Fig. 2B). We found minimal differences in speed between the two confinement states (Fig. 2C). We also did not find a difference between the entering and exiting speeds (Fig. 2D), nor between cells that had recently entered the channels and those that had been in the channels for a while (Fig. 2C,D).

**Figure 2.**
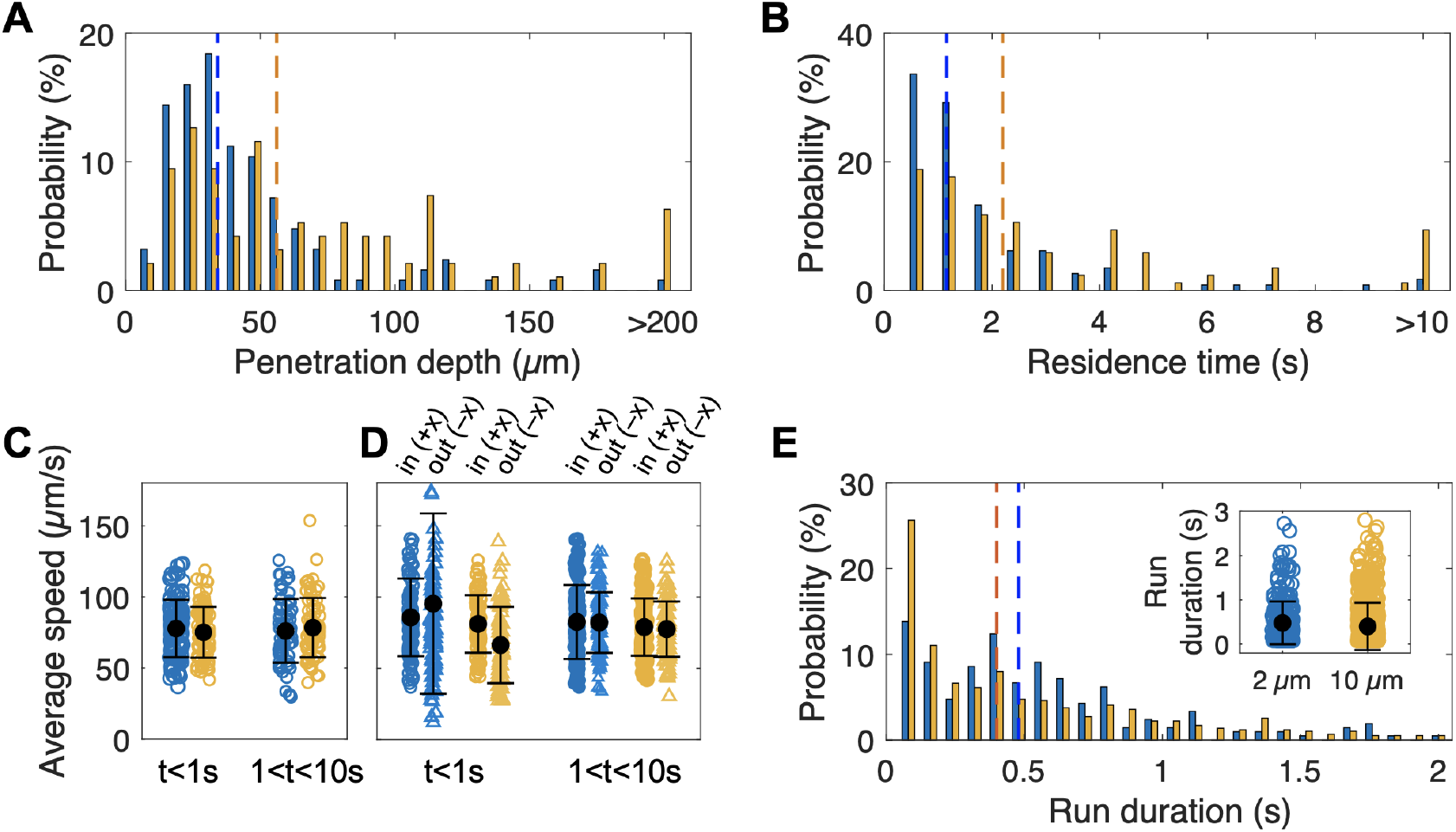
Confinement reduces penetrance and residence time of *V. fischeri*. A) Maximum penetration depth into channels of *V. fischeri* under 2 μm or 10 μm confinement. Dotted lines represent median. B) Channel persistence time of *V. fischeri* under 2 μm or 10 μm confinement. Dotted lines represent median. C) Average per cell swimming speed of *V. fischeri* during the t=0-1s or t=1-9s of each cell entering the confinement channel. Black dots and error bars=mean +/- SD. D) Average per cell speed of *V. fischeri* as it swims into (circles on left, positive x-direction) or out of (triangles on right, negative x-direction) confinement channels during the t=0-1s or t=1-9s intervals. Black dots and error bars=mean +/- SD. E) Histogram of run duration before a sharp (>120°) reversal of *V. fischeri* under 2 μm or 10 μm confinement. Dotted lines represent median. Inset show individual values, with black dots and bars representing mean +/- SD. For all panels, blue=2 μm, orange=10 μm.

### Confinement delays reversals

We next aimed to uncover what factors contributed to the stark divergence in swimming behavior as cells entered differently confined environments. One important distinction that emerged from comparing bacteria swimming in 2 μm and 10 μm channels is in the ‘run duration’, or the time interval between sharp reversals. The reversals were noted when the swimming direction changed by more than 120° within 80 ms. We observed that bacteria entering the 2 μm channels had a reversal pattern distinct from those entering 10 μm channels. Specifically, in the less-confined condition, there was an exponential decrease in the frequency distribution of run durations, indicating a random and equally distributed pattern of sharp reversals (31) (Fig. 2E). However, in the 2 μm channels, while quick reversals were relatively infrequent, there was a relative peak of reversals after ~0.5 s, followed by a random reversal frequency similar to those seen in 10 μm channels (Fig. 2E). These data indicate that confined cells display a more robust “refractory” period between reversals, during which they tend to have a period of longer swimming runs, both after entering the channels and between reversals.

### Confinement reduces directional deviation

In addition to a discrepancy in reversal patterns between cells in the 2 μm and 10 μm channels, we hypothesized that a cell rotating around the vertical z axis is expected to experience increased viscous drag in tighter gaps, resulting in a lower rotational diffusion coefficient. To measure this, we measured the bacterial swimming direction, which is predicted to change randomly over time due to thermal fluctuations in the surrounding fluid. Our analysis revealed that the bacteria undergo both small and large directional changes, the latter due primarily to sharp reversals. To estimate the rotational diffusion coefficient arising solely from thermal fluctuations, we removed sharp reversals (as defined earlier, directional changes exceeding 120° within 80 ms), and considered the mean of the square of the cumulative directional change over time (Fig. 3A). This analysis revealed that confined cells displayed lower changes in directional reorientation than unconfined cells (rotational diffusion coefficient [*D_r_*]: *D*_*r*,10μm_ = 3.8 rad^2^ s^−1^; *D*_*r*,2μm_ = 2.0 rad^2^ s^−1^). From these data we concluded that confined cells exhibited straighter swimming trajectories than their unconfined counterparts (Fig. 3A). We did not find evidence of either confinement-based orientation that could steer the bacteria towards an exit, or any association between angle positioning and swimming speed (Fig. S1A). Specifically, we observed no association between orientation angle and angle change (Fig. S1B) or between orientation angle and speed change (Fig. S1C) in either the confined or unconfined channels.

**Figure 3.**
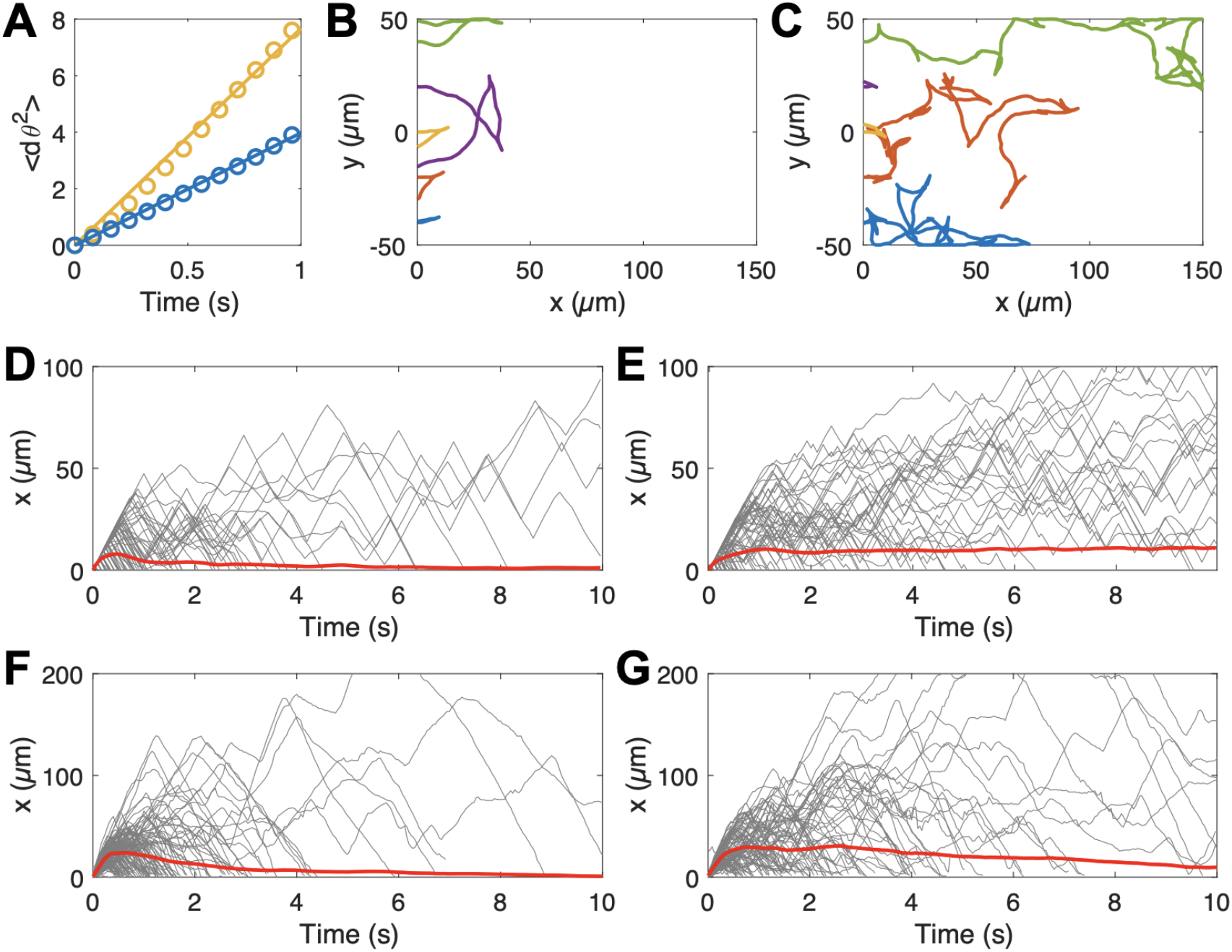
Confined cells have lower diffusional rotation, facilitating escape behavior. A) (delta-theta)^2^ over time of cells in 10 μm (orange) or 2 μm (blue) confined channels. The slope of this line approximates 2\**D_r_*. Circles represent experimental data, line is best linear fit. B) Simulation of cells swimming with a diffusional rotation constant (*D_r_*) of 0.5 rad^2^ s^−1^ and a 0.5 s delay between periods of random reversal, approximating activity seen in 2 μm channels. C) Simulation of cells swimming with a diffusional rotation constant (*D_r_*) of 1 rad^2^ s^−1^ and random reversal frequency, approximating activity seen in 10μm channels. D) Spatiotemporal plots of x-penetrance over time for cells in B. E) Spatiotemporal plots of x-penetrance over time for cells in C. F) Experimental spatiotemporal diagrams for *V. fischeri* swimming into 2 μm confinement chamber. G) Experimental spatiotemporal diagrams for *V. fischeri* swimming into 10 μm confinement chamber. In B and C, each line represents the trajectory of one simulated cell. In D-G, each black line represents the x position of one cell, and red lines represent the mean position of all cells that have entered the channel.

Motivated by the delayed reversals and the reduced rotational diffusion coefficient of bacteria in the relatively tighter confinement, we developed a simplified model to simulate the bacteria swimming with these differing traits (see Methods and SI). By varying *D_r_* and the delay in reversals while keeping all other parameters fixed, we found a drastic qualitative difference in swimming behavior resembling our experimental observations (Fig. 3B,C). These behavioral changes were sufficient to confer a difference in channel residence (Fig. 3D,E). When we compared these simulated results with those from our experimental findings, we found similar overall patterns of penetrance and escape (Fig. 3F,G), demonstrating that changes in rotational diffusivity and reversal frequency are sufficient to result in confinement-associated escape behavior.

### Confinement-based swimming can be counterbalanced by chemotactic cues

After demonstrating the confinement-avoidance swimming behavior, we sought to explore how this behavior influenced another known modulator of swimming behavior: chemotaxis. For these experiments, we used serine, which is a strong chemoattractant for *V. fischeri* (32). To allow us to simultaneously confine the cells while applying the attractant, we confined the bacteria with precise-diameter glass capillaries (see Methods) rather than dead-end channels. Suspensions of *V. fischeri* were loaded into the capillaries and, after minimizing background flow, we monitored the distribution of the cells inside capillaries of different terminal diameters after addition of the chemoattractant outside the capillary tip. In tightly confined capillaries (1 μm or 2 μm minimum diameter), cells avoided the most confined region (tip) of the capillary, mirroring what we saw in our dead-end channels (Movies S4-6). Even with the presence of chemoattractant in the external fluid, cells avoided the tip, rarely passing through these regions despite the physical clearance to do so (Fig. 4A,B). Conversely, cells experiencing less confinement readily left the capillaries and swam towards the chemoattractant outside the tip (Fig. 4C).

**Figure 4.**
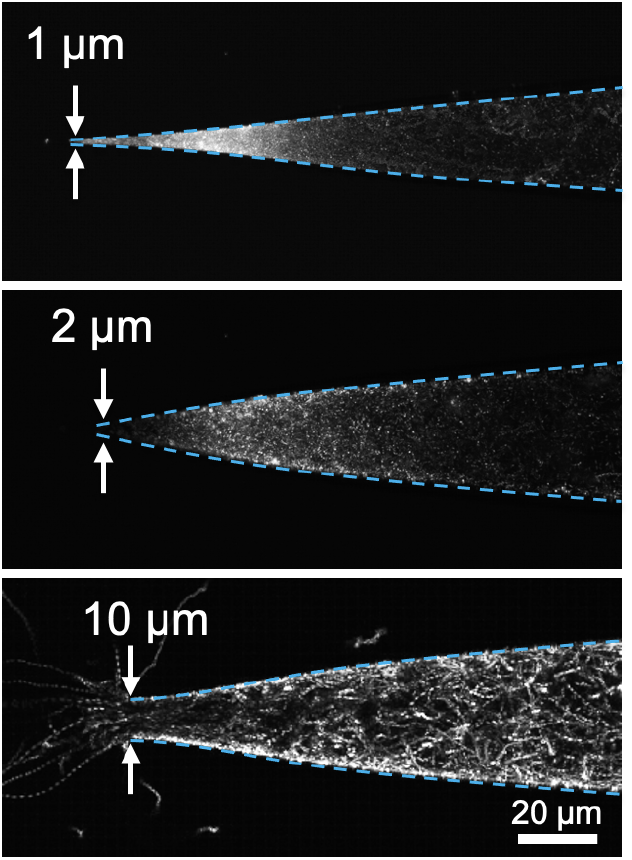
Chemoattraction counterbalances avoidance of confined spaces. Collapsed images from timelapse videos of GFP-labeled V. fischeri in glass capillaries with terminal diameter of 1 μm (top), 2 μm (middle), or 10 μm (bottom) after external addition of chemoattractant. Dotted cyan lines indicate edges of microcapillary.

## Discussion

The data presented here reveal that tight physical confinement can alter bacterial swimming behavior and long-term distribution by reducing directional fluctuations and delaying reversals. These experiments demonstrate that confinement interactions have profound implications on bacterial behavior, and may shape the patterns of motility during habitat transitions like those that occur during the initiation of symbiotic associations with animal and plant hosts.

The reversals we tracked under confined conditions could be the result of (i) the run-reverse activity used by many marine bacteria (33), (ii) the run-reverse-flick method employed by other *Vibrio* species (34), or (iii) flagellar wrapping, all of which allow quick, sharp-angle changes in swimming trajectories (11). Wrapping has been previously described in *Shewanella sp*., *Burkholderia sp*., and *V. fischeri* (10), and could result, at least partially, from interactions between the environment and the physical structure of the flagellar filament and its constituent flagellin proteins (35). The *V. fischeri* genome encodes at least five flagellin genes (36) and, as the time scale of reversal is too short to result from proteomic remodeling of these flagella (37), the specific individual combination of these flagellins could promote or inhibit the structural dynamics required for wrapping, potentially explaining heterogeneity among cells that quickly reversed rather than persisted in confinement. Also, our measurements showed that confined cells swam forward and backwards at similar speeds, whereas previous investigation of wrapping and run-reverse-flick behaviors has shown that these cells swim at different speeds when going forward and reverse (11, 38). Interestingly, a prior report has also described a difference in trajectory curvature between forward and reverse swimming, especially near a single surface (38). As we did not observe a noticeable difference in translational or angular speed by confined *V. fischeri* before or after reversals, two-sided confinement may balance out differences between reversal-associated swimming modes, potentially influencing reversals, escape behavior, and other directional movement (39–42).

The confined swimming behavior we observed may be selected for among bacterial species that experience confinement as a common part of their lifestyles. By preferentially reversing when confinement increases, cells could escape from places in which they might become stuck. There may even be a critical confinement point beyond which bacteria, especially rod- or comma-shaped bacteria like *V. fischeri*, may not be physically able to tumble or flick to facilitate an escape because of their shape and/or the length of their flagella. Such a constraint is exacerbated in bacteria like *V. fischeri* that can also produce extracellular capsule or polysaccharides, increasing their effective dimensions in certain environments (43, 44). These cells might avoid immobilization by quickly swimming away from tightly confined spaces and/or by wrapping their flagella to reduce their spatial footprint while swimming. In symbioses between flagellated bacteria and hosts, avoidance of being physically trapped could benefit both the bacterium and the host; if bacteria become trapped during colonization, they would no longer be able to swim towards favorable environments, while the host would experience clogging of bacteria-colonized tissues.

For bacteria like *V. fischeri*, which also must proceed through tightly confined spaces on the way to their target tissue, the balance between confinement avoidance and positive taxis (*e.g*., attraction to host-derived nutrients) may facilitate a stochastic procession through tightly confined spaces. This phenomenon may explain some of the extreme bottlenecking of *V. fischeri* cells on the way to the light organ crypts, which results in very few cells actually being able to colonize the symbiotic organ (45). Similar dynamics could play out in other symbiosis with tightly confining symbiotic tissues (46, 47), or with bacteria shaped like *V. fischeri*, including several pathogens (48). The combination of straighter swimming trajectories and increased escape activity could promote behavior where cells either make it through a tightly confined space to their target environment, or back out before they get stuck. This same principle could be applied to sort or separate cells based on their motility; physically confined environments could be experimentally shaped to selectively enrich for or against cells with this type of confinement-associated swimming behavior. Previously, similar physical heterogeneity has been shown to powerfully guide bacterial sorting (26). Our findings could inform the creation of apparatuses that spatially organize swimming cells by directing their confinement, providing a simple means to shape bacterial populations and communities. Our results also open the door for further exploration of the physical and biological determinants of confined swimming, holding major implications for fields such as host-microbe symbiosis and microbial biophysics.

Our findings open new questions about how bacteria and other flagellated cells navigate their physical environments, and how they modify their behaviors according to their surroundings. Questions remain about how different bacteria respond to confinement, the physical and biological factors that promote or reduce this escape behavior, and the downstream consequences of swimming modulation in response to confinement. These findings could explain complex interactions between bacteria and their environments, providing insights into diverse fields such as biofouling and multispecies symbiosis.

## Materials and Methods

### Bacterial preparation

*Vibrio fischeri* wild-type strain ES114 (28) was grown in LBS medium (pH 7.5, 10 g L^−1^ tryptone, 5 g L^−1^ yeast extract, 342 mM NaCl, 20 mM Tris (49)) at 28 °C. To label cells with GFP, plasmid pVSV102 (Kan^r^, (50)) was conjugated into *V. fischeri* using a previously described protocol (51). GFP-expressing cells were isolated on LBS plates containing kanomycin (50 μg mL^−1^) and stored in 25% glycerol at −80 °C.

For experimental conditions, individual colonies of GFP-labelled *V. fischeri* were placed in 1 mL of SWT medium (70% ocean water, 5 g L^−1^ tryptone, 3 g L^−1^ yeast extract, 32.5 mM glycerol (51)) for 90 minutes at 28 °C; these cells were pelleted at 8000 x g for 5 min. The supernatant was removed, and the cells were resuspended in 1 mL of filter-sterilized ocean water (FSOW), and left at room temperature throughout imaging.

For chemoattraction experiments, L-serine was dissolved in FSOW to a final concentration of 40 mM, and added outside of the end of a capillary tube.

### Fluorescence open space imaging

*V. fischeri* ES114 was subcultured from a single colony as described above, pelleted, and resuspended in FSOW, then stained using FM-464FX (ThermoFisher, Waltham, MA) according to previous protocols (10). About 10 μL of cell suspension was spotted inside of a fingernail polish ridge forming a square on a clean glass slide, and was sealed under a glass coverslip. Cells were imaged on a Nikon Eclipse Ti TIRF microscope using a 100X NA 1.5 TIRF objective (Nikon, Melville, NY) at 33 frames s^−1^ rate with a Cascade 512B EMCCD camera (Photometrics, Tucson, AZ). FM4-64 was excited at 532 nm (~ 10% laser power) and the emission was spectrally filtered (ZET532-10/ZT594rdc/ET645-75m; Chroma, Bellows Falls, VT) within the infinity space prior to reaching the detector.

### Motility measurements

Bacterial motility experiments were performed in two different settings: (i) dead-end microfluidic channels, and (ii) pulled glass capillaries. For channel experiments, a microfluidics device made of polydimethylsiloxane (PDMS) contained a straight-flow channel with width x height of 200 μm x 100 μm. At the middle of the channel, an array of narrow dead-end pores is laid perpendicular to the flow channel (see (27) for example). The height of the dead-end channels was either 2 μm or 10 μm, while the width and the length of the channels were kept at 45 μm and 400 μm, respectively. The motion of bacteria was observed using an inverted optical microscope (IX73, Olympus, Center Valley, PA) and a high-speed camera (Phantom Miro M110, Vision Research, Wayne, NJ). For the capillary experiments, pulled glass microcapillaries (World Precision Instruments, Sarasota, FL) with a tip-opening diameter of between 1 μm and 10 μm were placed in a closed chamber that was filled with filtered-sterilized seawater. The other end of the capillary was connected to a suspension of *V. fischeri*, which is placed on a vertical stage. By controlling the height of the reservoir, the background fluid flow can be mitigated. A small drop of 40 mM serine (32) was placed in the chamber at the tip of the capillary, and the motion of bacteria was observed using an inverted fluorescent microscope (DMi8, Leica, Buffalo Grove, IL).

### Image processing and data analysis

Cellular positioning was manually tracked at a rate of 25 frames s^−1^ by three independent researchers using Tracker 5.1.5 (https://physlets.org/tracker/). The x-y coordinates were exported and used to calculate position, speed, and angle with MATLAB (Mathworks, Portola Valley, CA).

### Computational modeling

We performed simulations of a dilute suspension of bacteria swimming independently in two dimensions. Each bacterium was assumed to swim at a constant speed (U = 50 μm s^−1^) in a direction that changes randomly due to thermal fluctuations and reversals. The reversals are modeled as a stochastic process such that after a fixed refractory period of either 0 or 0.5 s when no reversal occurs, the next reversal occurs randomly with mean rate f = 3 s^−1^and independently of the time since the refractory period. Otherwise, the direction changes continuously with rotational diffusion coefficient *D_r_* = 0.5 rad^2^ s^−1^ or 1.0 rad^2^ s^−1^. The resultant trajectories and directional changes were simulated In MATLAB (see SI for further details).

## Supporting information

Movie Supp. 1. Open swimming

Movie Supp. 3. Confined channels

Movie Supp. 5. 2um capillary

## Acknowledgments

This work was funded by NIH grant F32 GM119238 and a Ford Foundation Postdoctoral Fellowship to J.B.L., a C-MĀIKI seed grant from the University of Hawai‘i at Manoa to J.B.L and D.T., and NIH grant R01 GM135254 to E.G.R. and M.J.M.

---

**Figure S1.**
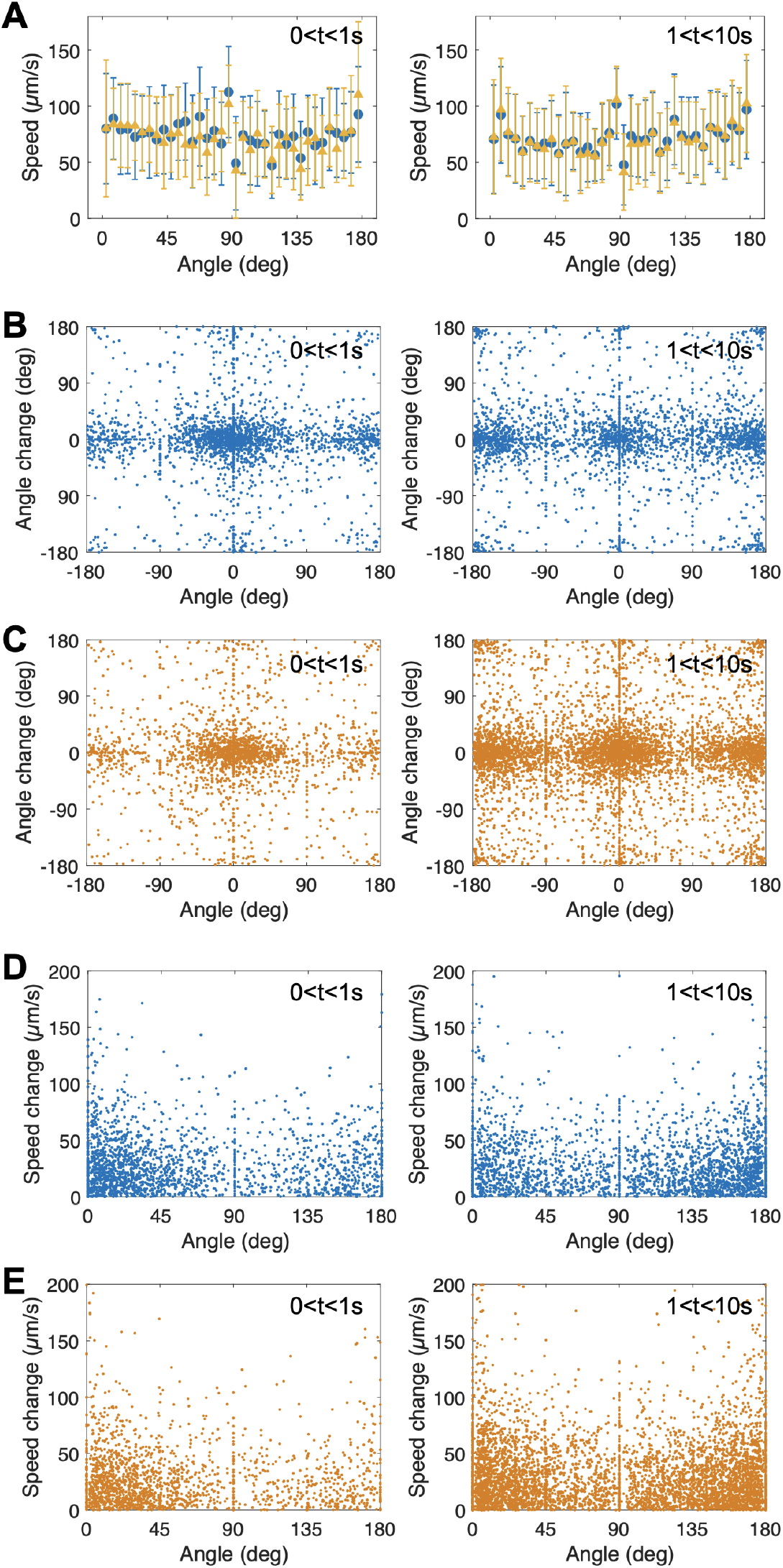
Angle change and speed are not correlated with orientation angle. Comparisons of A) orientation angle vs. absolute speed change, B,C) Angle change vs. orientation angle and D,E) orientation angle vs. speed. For each figure, data from cells in 2 μm confinement is blue and data from cells in 10 μm confinement is orange. Error bars=+/-SD.

Movie S1. Video of fluorescently labeled, unconfined *Vibrio fischeri* ES114 swimming near a surface. Cells were membrane-labeled with FM4-64 and imaged near the coverslip (see Methods) at a rate of 33 frames s^−1^.

Movie S2. Video corresponding to Figure 1B, where *V. fischeri* are swimming in a microfluidic device consisting of multiple less confined dead-end channels (height = 10 μm) connected to a large chamber (height = 100 μm).

Movie S3. Video corresponding to Figure 1C with similar experimental conditions as in Movie S2, but in more tightly confined dead-end channels (height = 2 μm).

Movie S4. Video corresponding to Figure 4, upper panel. *V. fischeri* are initially placed in a tapered capillary (tip diameter = 1 μm). Chemoattractants are subsequently released outside the capillary, allowing chemotaxis toward the tip.

Movie S5. Video corresponding to Figure 4, middle panel. Similar experimental conditions as in Movie S4, but with tip diameter of 2 μm.

Movie S6. Video corresponding to Figure 4, lower panel. Similar experimental conditions as in Movie S4, but with tip diameter of 10 μm.

