## Supplemental text file for "Transitioning to confined spaces impacts bacterial swimming and escape response"

PNAS
www.pnas.org

### Supplementary Information for

#### Transitioning to confined spaces impacts bacterial swimming and escape response

**Jonathan B. Lynch, Nicholas James, Margaret McFall-Ngai, Edward G. Ruby, Sangwoo Shin and Daisuke Takagi.**

**Jonathan B. Lynch.**
****

##### This PDF file includes:

Supplementary text
Table S1 (not allowed for Brief Reports)

#### Supporting Information Text

#### Computational modeling

**Two-dimensional space.** We consider a dilute suspension of bacteria moving horizontally in two dimensions. Minor vertical displacements in the third dimension were neglected because (i) the vertical displacements were not tracked in our experiments; and (ii) the channels constrained the bacteria to move primarily in the horizontal directions, especially in the channels with 2 $\mu$m spacing between the ceiling and the floor. The vast majority of bacteria remained far away from the side walls and the ends of the channels. Thus, they were neglected in the model.

**Stochastic reversals.** The trajectories of *V. fischeri* are characterized by 'run-and-reverse' dynamics consisting of relatively straight runs intervened by sudden reversals in swimming direction. The duration of each run is commonly assumed to have an exponentially decaying distribution, which arises if reversals occur randomly at some mean rate $f$ and independently of the time since the last reversal. However, our observations showed that bacteria in 2 $\mu$m channels have smaller occurrences of short runs (figure 2E), suggesting that some refractory period is needed after each reversal before the next reversal can occur. This motivated us to introduce a refractory period with no reversal for a fixed duration $T$, which was set to 0 or 0.5 s in 10 $\mu$m or 2 $\mu$m channels, respectively. Following this refractory period, the next reversal was assumed to occur randomly with mean rate $f = 3$ s$^{-1}$. The reversals, which consisted of changes in swimming direction by a distribution of angles ranging from approximately 120 to 180 degrees, were simplified with a fixed and instantaneous directional change of 180 degrees in our model.

**Thermal fluctuations.** Temporal changes in position and orientation of the bacteria were influenced significantly by thermal fluctuations. Our observations show that the swimming direction, which is expected to align approximately parallel to the long axis of the cell body, fluctuated considerably over time. By focusing on small angular displacements less than 120 degrees within 0.08 s to filter out the effects of reversals, and inspecting the linear growth in mean-square angular displacements with time due to thermal fluctuations, we estimated the rotational diffusion coefficient to be approximately 3.8 s$^{-1}$ or 2.0 s$^{-1}$ in 10 $\mu$m or 2 $\mu$m channels, respectively. The actual coefficients are likely to be smaller because flagellar activity may have produced additional directional changes. Nevertheless, the rotational diffusion coefficient is expected to be smaller in 2 $\mu$m than 10 $\mu$m channels because rotating bacteria are resisted by increased viscous drag in tighter confinement. To study the effects of reduced rotational diffusion in tighter confinement, we considered bacteria with a characteristic rotational diffusion coefficient of $D_r = 1.0$ s$^{-1}$ and reduced it by a factor of 2 in our model. The model neglected the effects of translational diffusion on the bacteria's position, which changed more importantly due to swimming. The swimming speed, which fluctuated with time in our experiments, was kept fixed at $U = 50$ $\mu$m s$^{-1}$.

**Simulations.** We simulated the bacterial dynamics using a discrete-time stochastic model. The model predicts the position and orientation of each bacterium using three time-dependent variables $(X, Y, \Theta)$, where $X$ and $Y$ denote the coordinates along and across the channel, respectively, and $\Theta$ denotes the angle in radians between the instantaneous swimming direction and the axis along the channel. The positions and directions were marched forward in time using the set of Langevin equations

$$X(t + \Delta t) = X(t) + U \cos[\Theta(t)]\Delta t, \quad [1]$$

$$Y(t + \Delta t) = Y(t) + U \sin[\Theta(t)]\Delta t, \quad [2]$$

$$\Theta(t + \Delta t) = \Theta(t) + \sqrt{2D_r\Delta t}A(t) + \pi B(t), \quad [3]$$

where $\Delta t = 0.04$ s is the time step, and $A$ and $B$ are pseudorandom numbers generated in MATLAB. We set $A = 0$ for the duration in time $T + G$ after each reversal, where $T$ is the fixed refractory period, and $G$ is the variable time until the next reversal after the refractory period. The variable times $G$ have a geometric distribution with small probability $p = f\Delta t = 0.02$, which ensured that reversals occur with mean rate $f$ and independently of the time after the refractory period. At the time step of reversal, we set $A = 1$. Pseudorandom numbers $B$ have a standard normal distribution to account for thermal fluctuations. We simulated $N = 100$ bacteria, all starting with $X = Y = \Theta = 0$ at time $t = 0$, and terminating at time $t = 10$ s. Any bacterium with $X < 0$ was marked as escaped.

Bacteria in 2 $\mu$m and 10 $\mu$m channels were simulated separately by adjusting the parameters $D_r$ and $T$, keeping the other parameters $U$ and $f$ fixed. The parameter values used in the simulations are shown in Table S1. These values are comparable in order of magnitude to those estimated empirically using our experimental data.

**Table S1. Table of parameter values used in computer simulations.**

| Confinement gap [$\mu$m] | 2 | 10 |
|---|---|---|
| Rotational diffusion coefficient $D_r$ [1/s] | 0.5 | 1.0 |
| Refractory period after each reversal, $T$ [s] | 0.5 | 0.0 |
| Mean reversal rate after refractory period, $f$ [1/s] | 3 | 3 |
| Swimming speed $U$ [$\mu$m/s] | 50 | 50 |
